# Predicting amphibian intraspecific diversity with machine learning: Challenges and prospects for integrating traits, geography, and genetic data

**DOI:** 10.1101/2020.05.03.073049

**Authors:** Lisa N. Barrow, Emanuel Masiero da Fonseca, Coleen E. P. Thompson, Bryan C. Carstens

## Abstract

The growing availability of genetic datasets, in combination with machine learning frameworks, offer great potential to answer long-standing questions in ecology and evolution. One such question has intrigued population geneticists, biogeographers, and conservation biologists: What determines intraspecific genetic diversity? This question is challenging to answer because many factors may influence genetic variation, including life history traits, historical influences, and geography, and the relative importance of these factors varies across taxonomic and geographic scales. Furthermore, interpreting the influence of numerous, potentially correlated variables is difficult with traditional statistical approaches. To address these challenges, we combined repurposed data with machine learning and investigated predictors of genetic diversity, focusing on Nearctic amphibians as a case study. We aggregated species traits, range characteristics, and >42,000 genetic sequences for 299 species using open-access scripts and various databases. After identifying important predictors of nucleotide diversity with random forest regression, we conducted follow-up analyses to examine the roles of phylogenetic history, geography, and demographic processes on intraspecific diversity. Although life history traits were not important predictors for this dataset, we found significant phylogenetic signal in genetic diversity within amphibians. We also found that salamander species at northern latitudes contain lower genetic diversity. Data repurposing and machine learning provide valuable tools for detecting patterns with relevance for conservation, but concerted efforts are needed to compile meaningful datasets with greater utility for understanding global biodiversity.

## Introduction

Genetic variation within species is a fundamental aspect of diversity that has long intrigued population geneticists, biogeographers, and conservation biologists (Avise, 2000; Lande, 1988; Leffler et al., 2012). Within species, genetic diversity includes both diversity within populations and genetic differences among populations, typically described as population genetic structure. Species with sufficient genetic diversity may have the ability to develop resistance to disease or adapt to environmental change (Ekroth, Rafaluk-Mohr, & King, 2019; Frankham, 1995; Jamieson & Allendorf, 2012). In contrast, species with limited genetic diversity can experience detrimental effects due to inbreeding depression and may be at higher risk of extinction from catastrophic events such as disease outbreaks (Frankham, 2005; Hedrick & Kalinowski, 2000; Lande, 1988). While the importance of conserving intraspecific diversity is well-established (Moritz, 2002; Moritz & Faith, 1998; Paz-Vinas et al., 2018), it remains to be fully integrated as standard practice in global conservation strategies (Laikre et al., 2020). In the past few decades, the availability of genetic datasets has grown dramatically, and large amounts of genetic data are now available to investigate diversity within species (Garrick et al., 2015). Furthermore, modern researchers have access to troves of data including species traits, genetic information, and geographic distributions from museum collections (e.g., <u>iDigBio</u>), genetic data repositories (e.g., <u>NCBI GenBank</u>), biodiversity portals (e.g., <u>GBIF</u>) and phenotypic databases (e.g., <u>MorphoBank</u>). Predicting and understanding the determinants of intraspecific genetic diversity broadly across taxonomic groups is an increasingly attainable goal.

Many potential predictors of intraspecific genetic diversity have been implicated. Effective population size and mutation rate are two key determinants of genetic diversity, each of which is governed by life history traits, which are expected to influence the accumulation and maintenance of genetic variation (Ellegren & Galtier, 2016). For example, Romiguier et al., (2014) investigated genome-wide diversity of 76 animal species and found a correlation between genetic diversity and life history strategy; long-lived species with few, large offspring and thus higher parental investment (“K-strategists”) had lower genetic diversity than species with many, small offspring (“r-strategists”). Chen, Glémin, & Lascoux, (2017) confirmed these results for animals and also found that longevity and mating system explained genomic diversity in plants. Body size has been negatively correlated with genetic diversity in various taxonomic groups including mammals (Brüniche-Olsen, Kellner, Anderson, & DeWoody, 2018), bees (López-Uribe, Jha, & Soro, 2019), and butterflies (Mackintosh et al., 2019), a relationship that is presumed to correspond with limits to population abundance in larger-bodied species (White, Ernest, Kerkhoff, & Enquist, 2007). Ecological traits associated with dispersal propensity, including body size, larval period, philopatry, or habitat specialization, could also influence whether or not species exhibit genetic structure (Mims, Hauser, Goldberg, & Olden, 2016; Paz, Ibáñez, Lips, & Crawford, 2015; Semlitsch, 2008). However, empirical support for associations between species traits and genetic diversity across taxonomic groups has been mixed. Within butterflies, Mackintosh et al., (2019) found no relationship between genetic diversity and longevity, propagule size, abundance, or host range. In Australian lizards, Grundler, Singhal, Cowan, & Rabosky, (2019) found no relationship between genetic diversity and body size or habitat specialization, and Singhal et al., (2017) found the only significant predictor of intraspecific diversity to be the number of museum occurrence records, a possible proxy for abundance. Investigations such as these exemplify the possibility that the determinants of genetic diversity may not be easily predictable and are likely to vary among taxonomic groups.

In addition to life history traits, geographic range characteristics can be useful metrics for predicting intraspecific genetic diversity. Total range size is sometimes treated as a proxy for census population size, which should scale positively with genetic diversity, because species that occupy more area or that have historically been able to expand to new areas are expected to have higher abundances (Leffler et al., 2012; Singhal et al., 2017). Additionally, total range size should be related to genetic structure because larger ranges typically encompass more habitat variation or physical barriers, leading to patterns of isolation by distance or environment, assuming the species are not capable dispersers (Sexton, Hangartner, & Hoffmann, 2014; Wright, 1943). Incorporating geographic variables such as elevational, topographic, or climatic variation has illustrated the influence of these factors on population connectivity and phylogeographic structure (Rodríguez et al., 2015; Wang, 2012). In a global study mapping amphibian genetic diversity, Miraldo et al., (2016) identified a latitudinal gradient of genetic diversity with higher diversity in the tropics. Similarly, at the intraspecific level, latitude has been negatively associated with genetic structure and diversity for a broad range of taxa (Brüniche-Olsen et al., 2018; Pelletier & Carstens, 2018; Smith, Seeholzer, Harvey, Cuervo, & Brumfield, 2017). A plausible explanation is that the climatic and environmental instability experienced by taxa at higher latitudes during glacial cycles contributed to this pattern by reducing suitable habitat and population sizes (Hewitt, 2000). The continued integration of intraspecific genetic variation, species traits, and geographic variables is needed to better understand the drivers of genetic diversity across different taxonomic groups and spatial scales.

Meta-analysis and data repurposing represent two strategies for integrating and analyzing complex data types from multiple species. Meta-analyses combine results from many studies to increase power and improve our understanding of key phenomena (e.g., Field et al., 2009; Soltis, Morris, McLachlan, Manos, & Soltis, 2006). Meta-analyses are able to draw from individual studies, each tailored to a focal taxon, but it is not easy to synthesize results across studies that differ in their aims, types of data, inference methods, and sampling design. It is often more efficient to proceed via data repurposing (Sidlauskas et al., 2010), where data from multiple studies are reanalyzed in a common framework. Repurposed data can be effectively combined with machine learning techniques to investigate questions of interest (e.g., Pelletier & Carstens, 2018). This strategy is consistent with broader goals in ecology and evolution to make data accessible and standardized (Blanchet, Prunier, & De Kort, 2017; Carpenter et al., 2009; Parr, Guralnick, Cellinese, & Page, 2012; Peters, 2010; Pope, Liggins, Keyse, Carvalho, & Riginos, 2015; Reichman, Jones, & Schildhauer, 2011; Sidlauskas et al., 2010; Soltis & Soltis, 2016). Furthermore, repurposing published data enhances their value and generates a greater return on investment in these data by funding agencies.

Of the many machine learning techniques available, we utilize the random forest approach introduced by Breiman (2001) because it is fast, easy to implement, can accommodate very large datasets including many predictor variables, and enables an intuitive evaluation of the importance of these variables for predicting the response. Random forest is a machine learning ensemble approach that uses multiple decision trees (a forest) to predict a user-defined response based on many potential predictor variables. Each individual decision tree consists of a subset of the data and a random subset of predictor variables at the nodes. Individually, each tree is a weak predictor, but when many trees are combined, random forest results in high predictive accuracy (Breiman, 2001). The importance of each variable is determined by examining the increase in prediction error after randomly permuting the data for that variable while all others are left unchanged. A review of random forest classification and regression is provided by (Liaw & Wiener, 2002). Here, we apply random forest regression to address a long-standing question in evolutionary biology (Leffler et al., 2012): What factors determine genetic diversity within species?

### Nearctic amphibians as a case study

Amphibians have long been considered the most imperiled class of vertebrates (Stuart et al., 2004). More than 40% of data-sufficient amphibian species are considered threatened by the International Union for Conservation of Nature (IUCN, 2019), and considerable efforts have focused on addressing the issue that a large proportion of species (∼25%) are data deficient, meaning there is insufficient information to assign them to an IUCN threat category (Nori, Villalobos, & Loyola, 2018). In a global analysis of amphibians, González-del-Pliego et al., (2019) found that body size and range size were correlated with threat status, estimated that half of data-deficient species are threatened, and identified regions of the world (e.g., the Neotropics and Southeast Asia) where a high number of data-deficient species are predicted to be threatened. Using machine learning, Howard & Bickford (2014) predicted that >63% of data-deficient species are threatened and identified similar geographic regions with high predicted extinction risk. However, potentially useful information is lacking from these studies. Notably, intraspecific genetic diversity and the factors that influence this measure are often absent in conservation assessments, despite their relevance to understanding the long-term persistence and adaptive capacity of species.

Our goal was to aggregate existing trait, geographic, and genetic data to investigate the predictors of genetic diversity in amphibians. We focused on Nearctic amphibians as an initial test case given the long history of study, tractable number of species, and presumed availability of natural history and genetic information. This dataset included all species native to Canada and the U.S., a subset of which have ranges extending into Mexico. First, we compiled species’ range variables, natural history characteristics, and gene alignments using a series of open-access databases and scripts. Second, we used random forest regression to identify predictors of intraspecific nucleotide diversity based on *Cytochrome b* (*cytb*), the gene with the most data available for amphibians and one that is commonly used for studying genetic variation within species (e.g., Vences, Thomas, Bonett, & Vieites, 2005). Finally, we investigated the relevance of phylogenetic history and demographic processes in contributing to current patterns of genetic diversity identified across amphibian species. We found evidence that genetic diversity within species is phylogenetically conserved, species traits do not predict genetic diversity for this set of species, and geographic range characteristics correspond with genetic diversity in Nearctic salamanders.

## Materials and Methods

### Species traits

We compiled trait data for amphibian species focusing on traits related to life history and ecology that may influence intraspecific genetic diversity. The following traits were initially extracted from the AmphiBIO database (Oliveira, São-Pedro, Santos-Barrera, Penone, & Costa, 2017): development mode (larval or direct), maximum body size (mm), age at maturity (min and max years), longevity (max years), clutch size (min and max number of eggs), and egg size (min and max in mm). We retrieved additional body size information from the Peterson field guides to reptiles and amphibians of the United States (Powell, Conant, & Collins, 2016; Stebbins & McGinnis, 2018). We used maximum snout-vent length (SVL) for Anura (frogs), and maximum total length for Caudata (salamanders; note that SVL was not always available for Caudata). The remaining traits of interest (neoteny, breeding habitat, larval period, time to hatching, dispersal distance, and home range size) were compiled primarily from species accounts in (AmphibiaWeb, 2018) <https://amphibiaweb.org>. Neoteny, or the retention of juvenile characteristics as adults, was categorized as ‘yes’ (a species is always neotenic), ‘no’ (neoteny has never been reported), or ‘some’ (neoteny has been reported in some populations, but not all). Breeding habitat, including the habitat in which eggs and/or larvae develop, was categorized as ‘terrestrial’, or ‘aquatic’, with aquatic species further categorized as ‘permanent’, ‘ephemeral’, or ‘aquatic generalist’. Larval period was first recorded numerically (min and max days) and then binned into categories: ‘none’ (direct developing), ‘short’ (0–90 days), ‘mid’ (91–365 days), or ‘long’ (>1 year). Time to hatching (min and max days), dispersal distance (max recorded in meters), and home range size (max recorded in meters squared) were recorded when available, but were unavailable for >40% of species and were thus excluded from further analyses in the present study (Fig. S1).

### Range characteristics

Species range maps for amphibians were downloaded as shapefiles from the IUCN Red List of Threatened Species Version 6.1 (www.iucnredlist.org/) on 22 January 2019. Range maps were manually edited in QGIS v. 2.18.2 as needed to exclude non-native ranges originally included in IUCN range maps, and to generate range maps for species that have been described recently from a portion of a former species’ range (e.g., *Acris blanchardi* split from *Acris crepitans*; Gamble, Berendzen, Bradley Shaffer, Starkey, & Simons, 2008). Revised shapefiles are available on Dryad (https://doi.org/10.5061/dryad.0cfxpnvzh; Barrow, Fonseca, Thompson, & Carstens, 2020). Altitude and bioclimatic variables were downloaded from WorldClim v. 1.4 (Hijmans, Cameron, Parra, Jones, & Jarvis, 2005) at a spatial resolution of 2.5 minutes (∼21 km^2^). We extracted the following information from the list of focal amphibian species: total range size (km^2^), latitude (min, max, extent = max-min, midpoint = (max+min)/2), altitude (min, max, average, standard deviation (SD)), slope (min, max, average, SD, extent), and bioclimatic variables (min, max, average, SD) using a custom script in R and the packages ‘rgdal’ (Bivand, Keitt, & Rowlingson, 2019), ‘raster’ (Hijmans, 2019b), and ‘geosphere’ (Hijmans, 2019a). Analyses were conducted in R v. 3.6 (R Core Team, 2019) and the scripts and datasets are available on Dryad.

### Genetic sequences

Sequences were downloaded from GenBank in two ways and were processed using a series of R and Python scripts modified from Pelletier & Carstens (2018). First, we obtained georeferenced sequences for all Nearctic amphibian species by querying accession numbers linked to records downloaded from (GBIF.org, 28 November 2018). Second, we downloaded all GenBank sequences for the 19 focal amphibian families, regardless of whether they were georeferenced, and only retained sequences from species within these families that have native ranges in the Nearctic. For both the georeferenced and non-georeferenced sequence sets separately, we sorted sequences by species and gene and generated multiple sequence alignments for every species by gene combination using Mafft v. 7.402 (Katoh & Standley, 2013). After summarizing the number of species and locus lengths for each species, we determined that cytochrome b (*cytb*) was the best-represented gene (Fig. S2). Alignments for *cytb* were then visually inspected in Geneious v. 8.1.9 (https://www.geneious.com) and edited to remove misaligned sequences prior to further analysis (<0.3% removed). Alignments for six additional anuran species were added from previously published datasets (Barrow, Bigelow, Phillips, & Lemmon, 2015; Barrow, Soto-Centeno, Warwick, Lemmon, & Moriarty Lemmon, 2017) available on Dryad or as unannotated mitochondrial genomes on GenBank (Accession numbers MF198257–MF198403; note these were missed by our scripts because gene names are not defined on GenBank, but sequences could be manually aligned to known *cytb* sequences).

For each species, we calculated nucleotide diversity using the nuc.div() function in the ‘pegas’ R package (Paradis, 2010). To assess sensitivity of nucleotide diversity to sample size, or the number of individuals sequenced, we randomly subsampled sequences from each *cytb* dataset with replacement to generate 100 datasets for each sample size between two and 25 sequences. We calculated nucleotide diversity for each dataset and examined the variance in the estimates from the 100 replicate datasets for each sample size. Given the steep decline in variance observed as sample size increased above five (Fig. S3), we conducted analyses of genetic diversity with species that included at least five sequences. Analyses were repeated using (1) the original value of nucleotide diversity for the full datasets and (2) the median value of nucleotide diversity from the 100 datasets with five randomly sampled sequences. Given the consistent results between the two datasets, we report results only for the latter. The R code and results for all analyses are available on Dryad.

### Random forest regression

We used random forest regression in the R package ‘randomForest’ (Liaw & Wiener, 2002) to predict intraspecific nucleotide diversity based on species traits (body size, development type, breeding habitat, neoteny, clutch size, and larval period), range characteristics (total range size, latitude, altitude, slope, and bioclimatic variables), and three additional predictors: taxonomic family, the number of GBIF occurrences, and the number of sequences for each species. We grew 2000 trees and used the defaults of randomly sampling one third of the predictors at each node (‘*m*_try_’) and minimum of five terminal nodes (‘nodesize’). Variable importance was determined based on the percent increase in mean squared error (IncMSE) of prediction when that predictor variable was randomly permuted (removing its effect) while all others were held constant. Model performance was assessed as the percent of variance in the response explained by the models. We conducted analyses for the dataset including species with at least five sequences and then conducted separate analyses for Anura and Caudata because some species traits are not comparable or applicable to both orders. For example, body size is measured differently between the two orders, only salamanders exhibit neoteny, and very few frog species native to the region have direct development. To examine the consistency of variable importance, we conducted 100 iterations with 2000 trees each, then averaged the percent variance explained and IncMSE for each variable across iterations. After conducting analyses with all variables of interest, we repeated analyses with reduced sets of predictors for ease of visualizing and interpreting the results.

### Phylogenetic comparative methods

Random forest analyses pointed to the importance of taxonomy for predicting nucleotide diversity (see Results). To better understand these results, we downloaded phylogeny subsets from VertLife.org for the Nearctic amphibian species in our dataset. Phylogenetic relationships are from Jetz & Pyron (2018) and are based on a time-calibrated posterior tree distribution for nearly all extant amphibian species, generated using a combination of phylogenetic inference and taxonomic assignment. We tested for phylogenetic signal in continuous life history traits and nucleotide diversity using the phylosig() function and one tree randomly sampled from the posterior. To visualize the differences in genetic diversity among species, we mapped median *cytb* nucleotide diversity onto the tree as a continuous trait using the contMap() function in the R package ‘phytools’ (Revell, 2012). This function estimates maximum likelihood ancestral traits at internal nodes and interpolates the states along the branches based on Felsenstein (1985). Additional analyses and tree pruning were conducted using the R package ‘ape’ (Paradis & Schliep, 2019). To account for phylogenetic relatedness between important continuous variables (see Results), we calculated phylogenetically independent contrasts using the pic() function, which implements the method described by Felsenstein (1985). We then assessed relationships between these continuous variables using linear models.

### Species distribution models

Random forest analyses indicate that the minimum latitude of a species was an important predictor of genetic diversity (see Results). We hypothesized that the signature of past demographic events, including population bottlenecks or post-glacial population expansion, would be more pronounced in species at higher (more northern) latitudes. We predicted that: (1) species at northern latitudes would have been more likely to experience range contraction during the last glacial maximum (LGM) than species at southern latitudes, and (2) species at northern latitudes would have low niche stability, or minimal overlap between the LGM and present suitability models. To test these predictions independent of the genetic data, we modeled species distributions based on current climate and hindcasted these to the LGM climate conditions. Occurrence records for each species were downloaded from (GBIF.org, 04 July 2019) using a polygon to encompass the ranges of all Nearctic amphibians. Custom scripts (available from Dryad, Barrow et al., 2020) were used to generate a file of occurrence localities for each species with *cytb* data, removing any points that fell outside the IUCN range map (0.5 degree buffer width) for that species, and thinning occurrence points to retain unique localities at a threshold of 10 km using the R package ‘spThin’ (Aiello-Lammens, Boria, Radosavljevic, Vilela, & Anderson, 2015). For species with a very large number of occurrences, we randomly sampled 3,000 localities prior to thinning to save computational time, and for species with small ranges and <25 localities, we did not thin datasets. We estimated species distribution models (SDMs) for 138 species with *cytb* data.

We downloaded 19 bioclimatic layers from WorldClim v. 1.4 (Hijmans et al. 2005) for current climate at a resolution of 30 seconds (∼1 km^2^) and for the LGM (CCSM4) at the highest resolution available (2.5 min). We conducted SDM analyses in R using the packages ‘biomod2’ (Thuiller, Georges, Engler, & Breiner, 2019), ‘raster’ (Hijmans, 2019b), ‘rgdal’ (Bivand, Keitt, & Rowlingson, 2019), ‘rgeos’ (Bivand & Rundel, 2019), and ‘HDInterval’ (Meredith & Kruschke, 2018). Bioclimatic layers for each time period were stacked, clipped to the extent of North America (−170, -52, 12, 72), and masked by current land borders. We tested for correlations between current climate layers and removed highly correlated variables (>0.8; Peterson, 2011), retaining five temperature (bio2, bio3, bio7, bio8, and bio10) and five precipitation variables (bio13, bio14, bio15, bio18, and bio19). Ensemble models were generated with the Generalized Linear Models (‘GLM’), Random Forest (‘RF’), and Maxent (‘MAXENT.phillips’) modeling options in ‘biomod2’. We used five model replicates for cross-validation, each with 80% of the data for training and 20% for testing and evaluated model accuracy with the ‘ROC’ evaluation metric. Models were hindcasted onto the LGM climate and were projected onto the current climate with the same resolution as the LGM (2.5 min).

For each time period, the five replicate models were averaged prior to calculating niche stability metrics. We converted models to binary presence/absence layers using a 95% occurrence probability threshold. The resulting models include the total area in which the model probability is equal to or greater than the probability threshold where 95% of the occurrence points are contained. For each threshold, we calculated the total suitable area for current and LGM models and determined whether the suitable area in the LGM was smaller (range contraction in the LGM) or larger (range expansion in the LGM) than the suitable area in the current model. We also determined the area of overlap between the two time periods and calculated the overlap index (OI) metric described by Hijmans & Graham (2006). The OI describes the proportion of the current range that was also suitable during the LGM, with higher values indicating higher niche stability. We compared the minimum latitude of species that were (1) predicted to undergo range contraction versus expansion and (2) species that had no niche overlap versus some niche overlap using Student’s t-tests in R.

## Results

### Data summary

We compiled data for 299 amphibian species native to the U.S. and Canada, 44 of which have ranges extending into Mexico. The dataset consists of 197 salamanders in nine families and 102 frogs in 10 families. Complete trait data were available for body size, development, breeding habitat, and neoteny, with various degrees of missing data for the remaining traits (Fig. S1). Range characteristics derived from IUCN range maps were available for 268 species, with data unavailable for recently described species (since 2008) or those with very small ranges. We assembled 4,759 georeferenced GenBank sequences that were linked to GBIF records, but only 37 species (20 salamanders, 17 frogs) were represented by at least five sequences from at least two localities. Preliminary analyses indicated there was no predictive power in models of genetic structure with this small dataset. We therefore focused analyses on non-spatial genetic datasets and used metrics of overall intraspecific genetic diversity. We assembled 42,067 GenBank sequences from 263 species, which consisted primarily of mitochondrial genes (Fig. S2). The best represented gene was *cytb*, with at least five sequences available for each of 147 species.

### Predictors of intraspecific genetic diversity

Full models with all predictors included 137 amphibian species with *cytb* data. These models explained an average of 24.9% variance in median nucleotide diversity based on 100 sequence datasets with five randomly sampled sequences. The most important predictors were consistently taxonomic family and the number of sequences, indicating that nucleotide diversity is influenced by phylogenetic relatedness and sample size. Some bioclimatic variables, including the average precipitation of the wettest month (bio13), average precipitation of the warmest quarter (bio18), average precipitation of the driest quarter (bio17), and the standard deviation of mean temperature of the driest quarter (bio9), were also ranked as important. Models with a subset of predictors explained an average of 30.1% variance in median nucleotide diversity. Taxonomic family, the number of sequences, and average precipitation of the wettest month (bio13) remained the most important predictors of nucleotide diversity, while species traits were not predictive of nucleotide diversity for this set of species (Fig. 1a).

**Figure 1.**
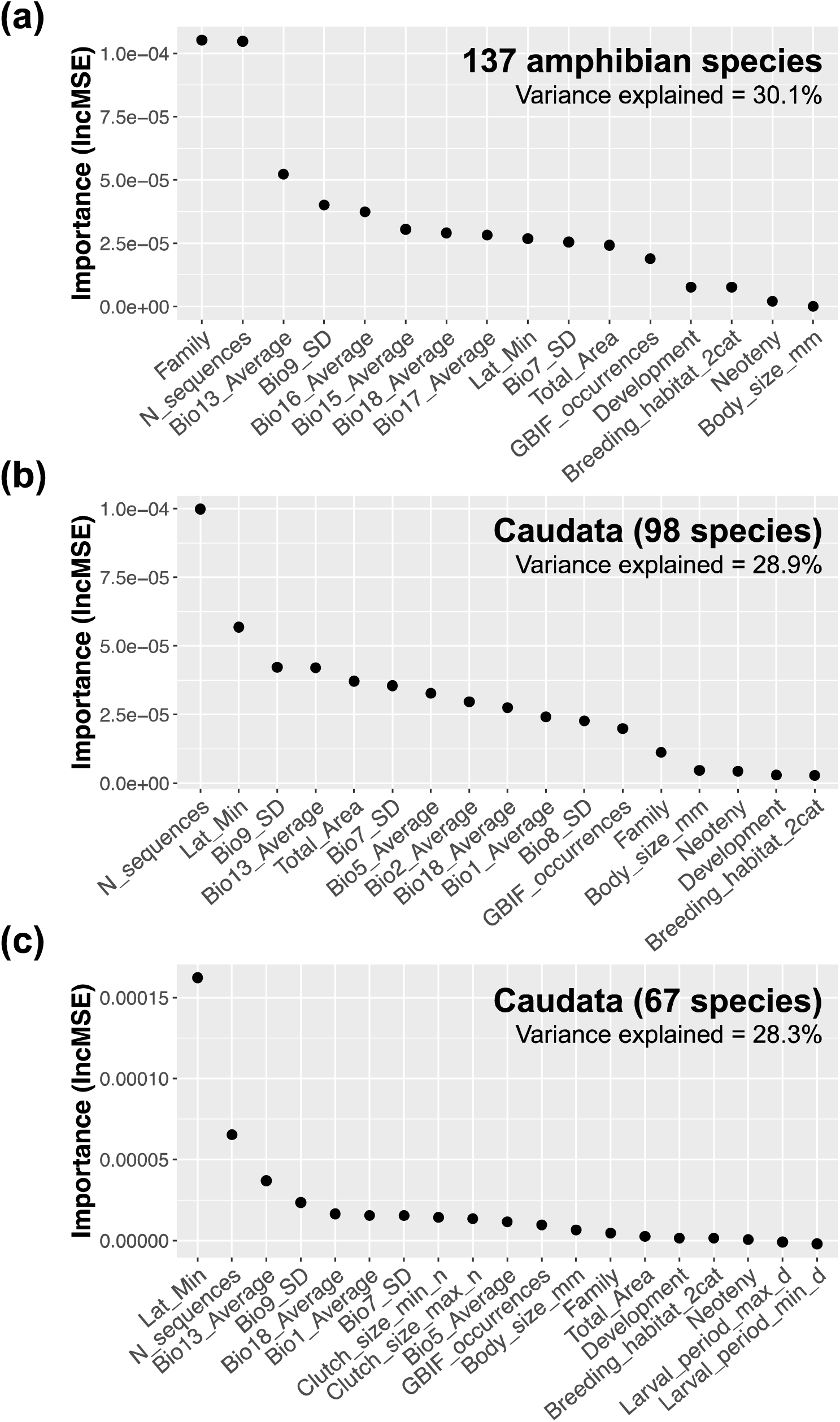
Predictors of intraspecific *cytb* nucleotide diversity for a) 137 species of Nearctic amphibians, b) 98 species of Caudata, and c) 67 species of Caudata with additional species traits. Average variable importance (increase in mean squared error) for each predictor from 100 iterations is shown.

When we considered each order separately, the Anura dataset included only 39 species with *cytb* data and there was no predictive power in these models (variance explained was negative). Full models for Caudata, however, included 98 species and explained an average of 20.5% of the variance in median nucleotide diversity. The most important predictors of nucleotide diversity were consistently the number of sequences and the minimum latitude of a species range. Models with a subset of predictors explained an average of 28.9% of the variance in median nucleotide diversity. The number of sequences and minimum latitude were the most important predictors (Fig. 1b). Models with fewer species (n=67) and additional trait information available explained an average of 28.3% variance in median nucleotide diversity. Minimum latitude remained the most important predictor and species traits were not important predictors of nucleotide diversity (Fig. 1c).

### Phylogenetic signal, minimum latitude, and climatic niche stability

We found phylogenetic signal in species-wide nucleotide diversity (lambda = 0.152, p = 0.008). Phylogenetic signal is also present in several traits related to reproduction and ecology including clutch size, larval period, and body size (Table S1), although these traits do not predict nucleotide diversity (described above and further tested with phylogenetic generalized linear mixed models; results not shown). The salamander families Sirenidae and Plethodontidae included multiple species with high nucleotide diversity (Fig. 2), but within Caudata, phylogenetic signal in nucleotide diversity was not significant (lambda = 0.105, p = 0.605). Notably, there was no phylogenetic signal in the number of sequences sampled, suggesting that the phylogenetic signal present in nucleotide diversity is not an artefact of sampling.

**Figure 2.**
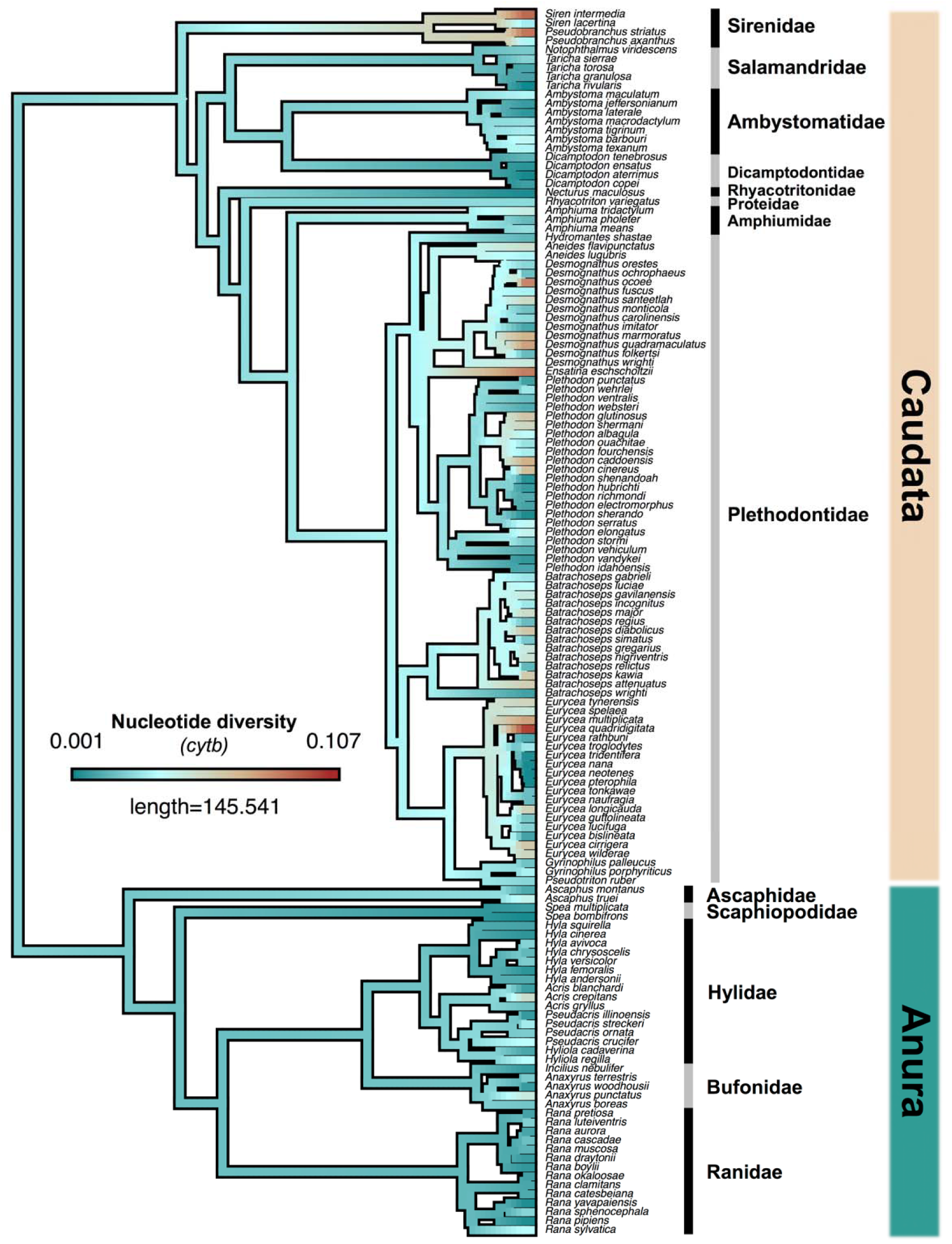
Phylogeny of Nearctic amphibian species with *cytb* nucleotide diversity mapped as a continuous trait. Phylogenetic relationships are based on Jetz and Pyron (2018) and were downloaded from VertLife.org, with length indicating time in millions of years. The median nucleotide diversity from 100 datasets with five randomly-sampled sequences was mapped onto the tree using the contMap() function on phytools (Revell, 2012). Warmer (brown/tan) colors indicate species with higher *cytb* nucleotide diversity.

After accounting for phylogenetic relationships there was a negative correlation between species-wide nucleotide diversity and minimum latitude in Caudata (Fig. 3). Species with more northern ranges tended to have lower nucleotide diversity (98-species dataset: R^2^ = 0.057, p = 0.0102; 67-species dataset: R^2^ = 0.115, p = 0.003). Plots of the raw values of nucleotide diversity and minimum latitude are shown in Fig. S4. Results from SDMs did not, however, support the hypothesis that species at higher latitudes were more heavily impacted by glaciation at the LGM. At the 95% occurrence probability threshold, we inferred LGM contraction of 76 Caudata species and LGM expansion for 20 species, but there was no difference in minimum latitude of the two groups (t = -1.59, p = 0.12). At the same threshold, 49 Caudata species had no overlap between the LGM and current models, indicating low niche stability, while 47 species had at least some overlap, indicating predicted areas of stability. There was no difference in minimum latitude of the two groups (t = 0.83, p = 0.41).

**Figure 3.**
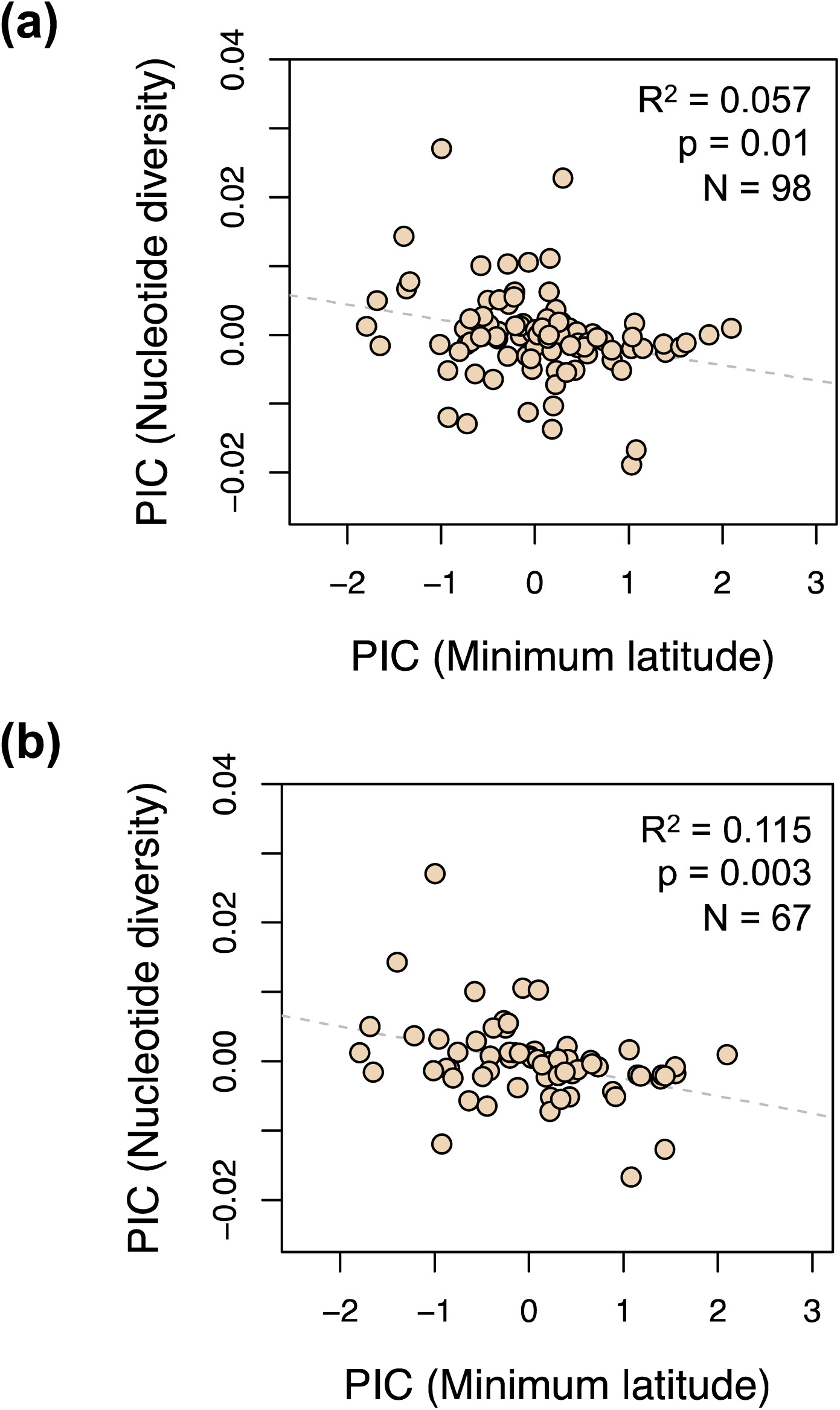
Minimum latitude and *cytb* nucleotide diversity for 98 Caudata species. a) Nucleotide diversity has a negative relationship with the minimum latitude of a species range. The median nucleotide diversity from 100 datasets with five randomly-sampled sequences is shown. b) Phylogenetic independent contrasts for the same metrics in a) using the pruned phylogeny from VertLife.org (Jetz and Pyron 2018).

## Discussion

Combining repurposed data with machine learning is a powerful strategy for addressing broad questions in evolutionary biology and conservation (Howard & Bickford, 2014; Sidlauskas et al., 2010). Here, we apply these approaches to investigate a classic question: What factors determine intraspecific genetic variation? We discuss results from our case study, including novel insights in Nearctic amphibian biogeography and broader implications that demonstrate the relevance of continued studies in this framework. We then highlight the prospects and challenges of combining data repurposing and machine learning to address questions that involve many complex data types and predictors.

### Predictors of genetic diversity in Nearctic amphibians

Amphibians play critical ecological roles as consumers (Whiles et al., 2006), prey (Zipkin, DiRenzo, Ray, Rossman, & Lips, 2020), and indicators of aquatic and terrestrial environmental health (Kerby, Richards-Hrdlicka, Storfer, & Skelly, 2010; Vitt, Caldwell, Wilbur, & Smith, 1990; Welsh & Ollivier, 1998). Given these roles, additional efforts to synthesize diverse datasets are warranted to inform conservation priorities. We found that minimum latitude predicted genetic diversity within salamanders, with generally lower intraspecific genetic diversity for species at higher (i.e., more northern) latitudes. Intraspecific genetic diversity also appears to be phylogenetically conserved, a finding perhaps explained by conserved life history traits such as clutch size or body size that are associated with intraspecific genetic diversity (e.g., (Paz et al., 2015; Romiguier et al., 2014). While life history traits are not predictors of genetic diversity for this dataset, taxonomic family was identified as important. This is likely driven by two salamander families, Sirenidae and Plethodontidae, which contain most species with the highest intraspecific diversity. Since more than half of the amphibian species in our dataset are plethodontid salamanders, the lack of a relationship between traits and genetic diversity might be explained by the limited trait variation (e.g., all are relatively small-bodied and have small clutch sizes) but considerable variation in genetic diversity within the group (Fig. 2).

The relevance of geography for understanding patterns of genetic diversity is well known (Hewitt, 2000; Wright, 1943). The negative association between latitude and intraspecific genetic diversity likely relates to population history since high intraspecific diversity at lower latitudes may be attributed to long-term stability (Miraldo et al., 2016) and low diversity at northern latitudes may be explained by population bottlenecks during glacial periods (Hewitt, 2000). Published phylogeographic studies on some of the species included in our study support this prediction, attributing lower genetic diversity at higher latitudes to postglacial range expansion (Carstens, Brunsfeld, Demboski, Good, & Sullivan, 2005; Highton & Webster, 1976; Radomski, Hantak, Brown, & Kuchta, 2020). However, the effect of past climate cycles on intraspecific genetic diversity is clearly dependent on refugial dynamics, as a species that is restricted into a single glacial refugium would exhibit different patterns than one where populations were isolated in separate refugia. One complicating factor that we did not address here is whether current species taxonomy is an appropriate unit of comparison, or whether some species include cryptic lineages that have not been formally described. Future studies with georeferenced localities and assignment of sequences to lineages to estimate within-population diversity should help clarify these issues further.

### Prospects and challenges for applying machine learning to repurposed data

Data repurposing can meet some of the same goals as meta-analysis, such as establishing what we (as a field) understand about key patterns and pinpointing underlying mechanisms(Carstens, Morales, Field, & Pelletier, 2018). Both types of investigation play an important role by potentially identifying connections among many variables. However, machine learning techniques make data repurposing more appealing than meta-analysis as a strategy for extracting the most information out of publicly available data. While meta-analyses certainly provide important insights, genetic datasets collected and analyzed separately across studies, which may use disparate sample designs and report incongruent metrics, can limit the number of useable datasets and make interpretation difficult (Emel & Storfer, 2012; López-Uribe et al., 2019). Once data are assembled, machine learning enables researchers to analyze very large datasets that incorporate many complex variables. By including a large suite of potential predictors, random forests can identify potential links among variables in an unbiased manner. This feature is useful for complex topics such as the determinants of genetic diversity, where researchers might be biased by previous findings and focus only on a subset of predictor variables. One caveat is that machine learning analysis is only as useful as the response variable used for a particular question. For example, while predictive models of binary questions (e.g., ‘LC’ versus ‘nonLC’ IUCN status; Fig. S5) will likely be able to explain more variation than those that rely on regression techniques, not all questions can easily be reduced to a binary outcome (e.g., a continuous response such as nucleotide diversity). In this case, the best random forest regression models explained ∼30% of the variance in nucleotide diversity, and the importance of the number of sequences per species points to sampling issues that may only be solved with the collection of additional data.

The application of machine learning techniques to repurposed data offers great potential for addressing key questions in ecology and evolutionary biology, but several challenges remain (Table 1). Our focus on assembling a curated set of species allowed us to identify several errors that would have been missed by automated pipelines. While useful for future work, this effort limited both the taxonomic and geographic scale of our study. For example, it seems plausible that the associations between life history traits and genetic diversity identified previously may only be apparent across broader taxonomic scales (e.g., Romiguier et al., 2014) or when more species can be included that span substantial trait variation (e.g., Brüniche-Olsen et al., 2018). Expanding the set of taxa from this study to a broader regional or global scale is a clear next step that will require addressing two substantial challenges. Compiling meaningful trait datasets for large numbers of species is time-consuming, and for many species, natural history information is not easily available. Solving this challenge will require collaborative efforts by the research community to standardize trait information, contribute to trait datasets, and make these datasets easily available (e.g., Oliveira et al., 2017). In the future, increasing efforts to digitize natural history collections and develop informatics tools will ultimately enable the incorporation of individual and population-level trait variation into these types of studies (Guralnick et al., 2016; Hedrick et al., 2020). The second major challenge to extending the scope of this study is the limited amount of georeferenced genetic data available for most species.

**Table 1.** Summary of challenges for repurposed data projects. Examples of challenges encountered in the present study are provided with potential avenues for correction in the future.

| Challenge | Example or Description | Potential for Correction |
| --- | --- | --- |
| Trait Data |  |  |
| Lack of natural history information | Difficult to measure traits such as dispersal, longevity, development time | • Promote and fund these efforts.<br>• Focus on traits that can be attained from natural history collections. |
| Recorded differently across sources | Body size for salamanders either SVL or TL depending on source | • Develop standards within a taxonomic group.<br>• Carefully check your data. |
| Time consuming, difficult to automate | Three of these authors spent weeks compiling data | • More collaborative efforts.<br>• Make trait tables open access and extendable by research community.<br>• Incorporate individual variation using museum specimens and open access databases. |
| Deciding how to treat quantitative traits | Min/max attainable for the most species, but may not capture variation of interest |  |
| Deciding how to treat categorical traits | Some ambiguity; variation across ranges for habitat type |  |
| Range Data |  |  |
| Taxonomy mismatches across databases | e.g., *Dryophytes/Hyla*, *Hyliola/Pseudacris* | • Check for synonyms manually.<br>• Use ‘taxize’ R package<sup>1</sup> |
| Species without IUCN range maps | Newly described or split species, e.g., *Acris blanchardi* | • Revise manually.<br>• Use occurrence records. |
| Introduced localities or errors in range maps or occurrence records | e.g., *Hyla cinerea* in Puerto Rico, *Rana pipiens* historical records throughout Mexico | • Check data; Revise range maps.<br>• Use script to remove records outside of range maps. |
| Genetic Data |  |  |
| Most sequences are not georeferenced | 4,759 GenBank accessions linked to GBIF localities of 42,067 assembled (<12%) | • Link sequences to georeferenced voucher specimens in databases.<br>• Include lat/lon when uploading. |
| Different genes sequenced, most mitochondrial | Fig. S1; 217 species represented by at least one gene; only 147 with cytb | • Standardize loci collected.<br>• Identify comparable metrics.<br>• Shift to genome-scale datasets. |
| Inconsistencies during GenBank upload | Different gene names (cytb, cytochrome b, Cyt-b, etc.) | • Use script to relabel names.<br>• Standardize names in databases. |
| Data not uploaded to GenBank correctly (or at all) | “UNVERIFIED” sequences without genes annotated properly were missed | • Manually add data from other sources (e.g., Dryad).<br>• Correct GenBank records. |
| Errors in automated alignments | Mis-aligned sequences identified, particularly when lengths differed | • Inspect alignments.<br>• Test different software and optimize settings. |
| Small or uneven sample sizes across species | N individuals sampled was associated with diversity | • Check sensitivity to sampling.<br>• Collect more data! |

<sup>1</sup>(Chamberlain & Szöcs, 2013). SVL = snout-vent length; TL = total length.

Two main axes of sampling effort are important to interpreting intraspecific genetic diversity: the number (and location) of individuals sampled and the number (and type) of loci sampled. Researchers have typically had to make a trade-off between these two for logistical reasons. Although the number of loci that can feasibly be sequenced is growing rapidly (Garrick et al., 2015), the vast majority of available sequences are not directly linked to localities (Marques, Maronna, & Collins, 2013). Increasing the amount of georeferenced genetic data available per species will enable within-population variation to be disentangled from among-population variation and will foster a better understanding of the mechanisms that influence intraspecific genetic diversity. Currently, the genetic sequences available for most animal species are mitochondrial genes and it remains unclear how representative mitochondrial DNA actually is for describing intraspecific genetic diversity (Bazin, Glémin, & Galtier, 2006; Galtier, Nabholz, GlÉmin, & Hurst, 2009; Mulligan, Kitchen, & Miyamoto, 2006). Furthermore, it is likely that available molecular markers do not accurately reflect adaptive variation (Mittell, Nakagawa, & Hadfield, 2015), which may be important for designing conservation strategies that promote species persistence (Moritz, 2002). Despite these issues, comparing genetic diversity among species yields important insights, and new findings will be enhanced by continued efforts to link existing genetic data to localities and to collect genome-scale data from a large number of non-model species encompassing variation in geographic complexity and life history traits.

## Conclusions

Identifying the predictors of intraspecific genetic diversity and the scale for which they are predictive are important goals for both evolutionary biology and applied conservation. We demonstrated the potential for combining repurposed data with machine learning techniques to predict genetic diversity in Nearctic amphibians. Our findings indicate that life history traits do not predict genetic diversity in this dataset, but future studies should incorporate additional species and trait variation on a global scale. Total range size is an important criterion for assessing species conservation risk, but it does not predict genetic diversity within Nearctic amphibians. Instead, we found that minimum latitude was an important predictor of genetic diversity within salamanders, suggesting that this aspect of a species’ range represents an important component to consider when assessing species risk. Northern latitude species that harbor low genetic diversity may be even more vulnerable to future climate change scenarios or disease outbreaks. Identifying these potential risks by combining new and existing datasets can lead to proactive management strategies that preserve remaining genetic diversity.

## Acknowledgements

We thank Drew Duckett, Flavia Mol Lanna, Danielle Parsons, Megan Smith, Tara Pelletier, Jamin Wieringa, and Tamaki Yuri for helpful discussions, and the many contributors to AmphibiaWeb, GenBank, GBIF, and the IUCN Red List for archiving the data that made this study possible. This work was supported by NSF DBI 1910623, The Ohio State University (OSU) President’s Postdoctoral Scholars Program, and the OSU Department of Evolution, Ecology and Organismal Biology. EMF thanks the Coordenação de Aperfeiçoamento de Pessoal de Nível Superior (CAPES) for his doctoral fellowship (process #88881.170016/2018).

## Data Accessibility

Scripts for data processing and analysis, DNA sequence alignments, and trait and climate datasets are available on Dryad doi: https://doi.org/10.5061/dryad.0cfxpnvzh (Barrow et al., 2020).

## Author Contributions

LNB and BCC conceived the ideas; LNB, EMF, and CEPT gathered and analyzed the data; LNB and BCC wrote the paper with input from all authors.

## References

Aiello-Lammens, M. E., Boria, R. A., Radosavljevic, A., Vilela, B., & Anderson, R. P. (2015). spThin: An R package for spatial thinning of species occurrence records for use in ecological niche models. Ecography, 38(5), 541–545. doi: 10.1111/ecog.01132

AmphibiaWeb. (2018). <https://amphibiaweb.org>. Retrieved December 18, 2018, from University of California, Berkeley, CA, USA. Accessed 18 Dec 2018. website: https://amphibiaweb.org

Avise, J. C. (2000). Phylogeography: the history and formation of species. Cambridge, Massachussetts: Harvard University Press.

Barrow, L. N., Bigelow, A. T., Phillips, C. A., & Lemmon, E. M. (2015). Phylogeographic inference using Bayesian model comparison across a fragmented chorus frog species complex. Molecular Ecology, 24(18), 4739–4758. doi: 10.1111/mec.13343

[dataset] Barrow, L. N., Fonseca, E. M., Thompson, C. E. P., & Carstens, B. C. (2020). Data from: Predicting amphibian intraspecific diversity with machine learning: Challenges and prospects for integrating traits, geography, and genetic data. Dryad Digital Repository, https://doi.org/10.5061/dryad.0cfxpnvzh.

Barrow, L. N., Soto-Centeno, J. A., Warwick, A. R., Lemmon, A. R., & Moriarty Lemmon, E. (2017). Evaluating hypotheses of expansion from refugia through comparative phylogeography of south-eastern Coastal Plain amphibians. Journal of Biogeography, 44, 2692–2705. doi: 10.1111/jbi.13069

Bazin, E., Glémin, S., & Galtier, N. (2006). Mitochondrial Genetic Diversity in Animals. Science, 312(April), 570–572.

Bivand, R., Keitt, T., & Rowlingson, B. (2019). rgdal: Bindings for the “Geospatial” Data Abstraction Library. R package version 1.4-8. (p. https://cran.r-project.org/package=rgdal.). p. https://cran.r-project.org/package=rgdal.

Bivand, R., & Rundel, C. (2019). rgeos: Interface to Geometry Engine - Open Source (’GEOS’). R package version 0.4-3. https://CRAN.R-project.org/package=rgeos.

Blanchet, S., Prunier, J. G., & De Kort, H. (2017). Time to Go Bigger: Emerging Patterns in Macrogenetics. Trends in Genetics, 33(9), 579–580. doi: 10.1016/j.tig.2017.06.007

Breiman, L. (2001). Random forests. Machine Learning, 45, 5–32. doi: 10.1201/9780367816377-11

Brüniche-Olsen, A., Kellner, K. F., Anderson, C. J., & DeWoody, J. A. (2018). Runs of homozygosity have utility in mammalian conservation and evolutionary studies. Conservation Genetics, 19(6), 1295–1307. doi: 10.1007/s10592-018-1099-y

Carpenter, S. R., Armbrust, E. V., Arzberger, P. W., Chapin, F. S., Elser, J. J., Hackett, E. J., … Zimmerman, A. S. (2009). Accelerate Synthesis in Ecology and Environmental Sciences. BioScience, 59(8), 699–701. doi: 10.1525/bio.2009.59.8.11

Carstens, B. C., Brunsfeld, S. J., Demboski, J. R., Good, J. M., & Sullivan, J. (2005). Investigating the evolutionary history of the Pacific Northwest mesic forest ecosystem: hypothesis testing within a comparative phylogeographic framework. Evolution, 59(8), 1639–1652. doi: 10.1111/j.0014-3820.2005.tb01815.x

Carstens, B. C., Morales, A. E., Field, K., & Pelletier, T. A. (2018). A global analysis of bats using automated comparative phylogeography uncovers a surprising impact of Pleistocene glaciation. Journal of Biogeography, 45(8), 1795–1805. doi: 10.1111/jbi.13382

Chamberlain, S. A., & Szöcs, E. (2013). taxize: taxonomic search and retrieval in R. F1000Research, 2(191).

Chen, J., Glémin, S., & Lascoux, M. (2017). Genetic diversity and the efficacy of purifying selection across plant and animal species. Molecular Biology and Evolution, 34(6), 1417–1428. doi: 10.1093/molbev/msx088

Ekroth, A. K. E., Rafaluk-Mohr, C., & King, K. C. (2019). Host genetic diversity limits parasite success beyond agricultural systems: A meta-analysis. Proceedings of the Royal Society B: Biological Sciences, 286(1911). doi: 10.1098/rspb.2019.1811

Ellegren, H., & Galtier, N. (2016). Determinants of genetic diversity. Nature Reviews Genetics, 17(7), 422–433. doi: 10.1038/nrg.2016.58

Emel, S. L., & Storfer, A. (2012). A decade of amphibian population genetic studies: Synthesis and recommendations. Conservation Genetics, 13(6), 1685–1689. doi: 10.1007/s10592-012-0407-1

Felsenstein, J. (1985). Phylogenies and the comparative method. American Naturalist, 125(1), 1–15.

Field, R., Hawkins, B. A., Cornell, H. V., Currie, D. J., Diniz-Filho, J. A. F., Guégan, J. F., … Turner, J. R. G. (2009). Spatial species-richness gradients across scales: A meta-analysis. Journal of Biogeography, 36(1), 132–147. doi: 10.1111/j.1365-2699.2008.01963.x

Frankham, R. (1995). Effective population size/adult population size ratios in wildlife: A review. Genetics Research, 66, 95–107. doi: 10.1017/S0016672308009695

Frankham, R. (2005). Genetics and extinction. Biological Conservation, 126(2), 131–140. doi: 10.1016/j.biocon.2005.05.002

Galtier, N., Nabholz, B., GlÉmin, S., & Hurst, G. D. D. (2009). Mitochondrial DNA as a marker of molecular diversity: A reappraisal. Molecular Ecology, 18(22), 4541–4550. doi: 10.1111/j.1365-294X.2009.04380.x

Gamble, T., Berendzen, P. B., Bradley Shaffer, H., Starkey, D. E., & Simons, A. M. (2008). Species limits and phylogeography of North American cricket frogs (Acris: Hylidae). Molecular Phylogenetics and Evolution, 48(1), 112–125. doi: 10.1016/j.ympev.2008.03.015

Garrick, R. C., Bonatelli, I. A. S., Hyseni, C., Morales, A., Pelletier, T. A., Perez, M. F., … Carstens, B. C. (2015). The evolution of phylogeographic datasets. Molecular Ecology, 24, 1164–1171. doi: 10.1111/mec.13108

GBIF.org (28 November 2018) GBIF Occurrence Download https://doi.org/10.15468/dl.r6bshz.

GBIF.org (04 July 2019) GBIF Occurrence Download https://doi.org/10.15468/dl.rk2srg.

González-del-Pliego, P., Freckleton, R. P., Edwards, D. P., Koo, M. S., Scheffers, B. R., Pyron, R. A., & Jetz, W. (2019). Phylogenetic and Trait-Based Prediction of Extinction Risk for Data-Deficient Amphibians. Current Biology, 29(9), 1557-1563.e3. doi: 10.1016/j.cub.2019.04.005

Grundler, M. R., Singhal, S., Cowan, M. A., & Rabosky, D. L. (2019). Is genomic diversity a useful proxy for census population size? Evidence from a species-rich community of desert lizards. Molecular Ecology. doi: 10.1111/mec.15042

Guralnick, R. P., Zermoglio, P. F., Wieczorek, J., LaFrance, R., Bloom, D., & Russell, L. (2016). The importance of digitized biocollections as a source of trait data and a new VertNet resource. Database, 2016. doi: 10.1093/database/baw158

Hedrick, B. P., Heberling, J. M., Meineke, E. K., Turner, K. G., Grassa, C. J., Park, D. S., … Davis, C. C. (2020). Digitization and the Future of Natural History Collections. BioScience, XX(X), 1–9. doi: 10.1093/biosci/biz163

Hedrick, P. W., & Kalinowski, S. T. (2000). Inbreeding depression in conservation biology. Annual Review of Ecology and Systematics, 31, 139–162.

Hewitt, G. (2000). The genetic legacy of the Quaternary ice ages. Nature, 405(6789), 907–913. doi: 10.1038/35016000

Highton, R., & Webster, T. P. (1976). Geographic protein variation and divergence in populations of the salamander Plethodon cinereus. Evolution, 30(1), 33–45.

Hijmans, R. J. (2019a). geosphere: Spherical Trigonometry. R package version 1.5-10. https://CRAN.Rproject.org/package=geosphere.

Hijmans, R. J. (2019b). raster: Geographic Data Analysis and Modeling. R package version 3.0-7. https://CRAN.R-project.org/package=raster.

Hijmans, R. J., Cameron, S. E., Parra, J. L., Jones, P. G., & Jarvis, A. (2005). Very high resolution interpolated climate surfaces for global land areas. International Journal of Climatology, 25(15), 1965–1978. doi: 10.1002/joc.1276

Hijmans, R. J., & Graham, C. H. (2006). The ability of climate envelope models to predict the effect of climate change on species distributions. Global Change Biology, 12(12), 2272–2281. doi: 10.1111/j.1365-2486.2006.01256.x

Howard, S. D., & Bickford, D. P. (2014). Amphibians over the edge: Silent extinction risk of Data Deficient species. Diversity and Distributions, 20(7), 837–846. doi: 10.1111/ddi.12218

IUCN. (2019). The IUCN Red List of Threatened Species. Retrieved March 6, 2020, from Version 2019-3 website: https://www.iucnredlist.org/resources/summary-statistics#Figure2

Jamieson, I. G., & Allendorf, F. W. (2012). How does the 50/500 rule apply to MVPs? Trends in Ecology and Evolution, 27(10), 578–584. doi: 10.1016/j.tree.2012.07.001

Jetz, W., & Pyron, R. A. (2018). The interplay of past diversification and evolutionary isolation with present imperilment across the amphibian tree of life. Nature Ecology and Evolution, 2(5), 850–858. doi: 10.1038/s41559-018-0515-5

Katoh, K., & Standley, D. M. (2013). MAFFT multiple sequence alignment software version 7: Improvements in performance and usability. Molecular Biology and Evolution, 30(4), 772–780. doi: 10.1093/molbev/mst010

Kerby, J. L., Richards-Hrdlicka, K. L., Storfer, A., & Skelly, D. K. (2010). An examination of amphibian sensitivity to environmental contaminants: Are amphibians poor canaries? Ecology Letters, 13(1), 60–67. doi: 10.1111/j.1461-0248.2009.01399.x

Laikre, L., Hoban, S., Bruford, M. W., Segelbacher, G., Allendorf, F. W., Gajardo, G., … Vernesi, C. (2020). Post-2020 goals overlook genetic diversity. Science, 367(6482), 1083–1085.

Lande, R. (1988). Genetics and demography in biological conservation. Science, 241(4872), 1455–1460. doi: 10.1126/science.3420403

Leffler, E. M., Bullaughey, K., Matute, D. R., Meyer, W. K., Ségurel, L., Venkat, A., … Przeworski, M. (2012). Revisiting an Old Riddle: What Determines Genetic Diversity Levels within Species? PLoS Biology, 10(9). doi: 10.1371/journal.pbio.1001388

Liaw, A., & Wiener, M. (2002). Classification and Regression by randomForest. R News, 2(3), 18–22.

López-Uribe, M. M., Jha, S., & Soro, A. (2019). A trait-based approach to predict population genetic structure in bees. Molecular Ecology, 28(8), 1919–1929. doi: 10.1111/mec.15028

Mackintosh, A., Laetsch, D. R., Hayward, A., Charlesworth, B., Waterfall, M., Vila, R., & Lohse, K. (2019). The determinants of genetic diversity in butterflies. Nature Communications, 10(1), 1–9. doi: 10.1038/s41467-019-11308-4

Marques, A. C., Maronna, M. M., & Collins, A. G. (2013). Putting GenBank Data on the Map. Science, 341(September), 1341.

Meredith, M., & Kruschke, J. (2018). HDInterval: Highest (Posterior) Density Intervals. R package version 0.2.0. https://CRAN.R-project.org/package=HDInterval.

Mims, M. C., Hauser, L., Goldberg, C. S., & Olden, J. D. (2016). Genetic Differentiation, Isolation-by-Distance, and Metapopulation Dynamics of the Arizona Treefrog (Hyla wrightorum) in an Isolated Portion of Its Range. 1–23. doi: 10.1371/journal.pone.0160655

Miraldo, A., Li, S., Borregaard, M. K., Flórez-Rodríguez, A., Gopalakrishnan, S., Rizvanovic, M., … Nogués-Bravo, D. (2016). An Anthropocene map of genetic diversity. Science, 353(6307), 1532–1535. doi: 10.1126/science.aaf4381

Mittell, E. A., Nakagawa, S., & Hadfield, J. D. (2015). Are molecular markers useful predictors of adaptive potential? Ecology Letters, 18(8), 772–778. doi: 10.1111/ele.12454

Moritz, C. (2002). Strategies to protect biological diversity and the evolutionary processes that sustain it. Systematic Biology, 51(2), 238–254.

Moritz, C., & Faith, D. P. (1998). Comparative phylogeography and the identification of genetically divergent areas for conservation. Molecular Ecology, 7(4), 419–429. doi: 10.1046/j.1365-294x.1998.00317.x

Mulligan, C. J., Kitchen, A., & Miyamoto, M. M. (2006). Comment on “Population size does not influence mitochondrial genetic diversity in animals.” Science, 314(5804), 1390. doi: 10.1126/science.1122033

Nori, J., Villalobos, F., & Loyola, R. (2018). Global priority areas for amphibian research. Journal of Biogeography, 45(11), 2588–2594. doi: 10.1111/jbi.13435

Oliveira, B. F., São-Pedro, V. A., Santos-Barrera, G., Penone, C., & Costa, G. C. (2017). AmphiBIO, a global database for amphibian ecological traits. Scientific Data, 4, 1–7. doi: 10.1038/sdata.2017.123

Paradis, E. (2010). pegas: an R package for population genetics with an integrated-modular approach. Bioinformatics, 26, 419–420.

Paradis, E., & Schliep, K. (2019). Ape 5.0: An environment for modern phylogenetics and evolutionary analyses in R. Bioinformatics, 35(3), 526–528. doi: 10.1093/bioinformatics/bty633

Parr, C. S., Guralnick, R., Cellinese, N., & Page, R. D. M. (2012). Evolutionary informatics: Unifying knowledge about the diversity of life. Trends in Ecology and Evolution, 27(2), 94–103. doi: 10.1016/j.tree.2011.11.001

Paz, A., Ibáñez, R., Lips, K. R., & Crawford, A. J. (2015). Testing the role of ecology and life history in structuring genetic variation across a landscape: A trait-based phylogeographic approach. Molecular Ecology, 24(14), 3723–3737. doi: 10.1111/mec.13275

Paz-Vinas, I., Loot, G., Hermoso, V., Veyssière, C., Poulet, N., Grenouillet, G., & Blanchet, S. (2018). Systematic conservation planning for intraspecific genetic diversity. Proceedings of the Royal Society B: Biological Sciences, 285(1877). doi: 10.1098/rspb.2017.2746

Pelletier, T. A., & Carstens, B. C. (2018). Geographical range size and latitude predict population genetic structure in a global survey. Biology Letters, 14(1). doi: 10.1098/rsbl.2017.0566

Peters, D. P. C. (2010). Accessible ecology: Synthesis of the long, deep, and broad. Trends in Ecology and Evolution, 25(10), 592–601. doi: 10.1016/j.tree.2010.07.005

Peterson, A. T. (2011). Ecological niche conservatism: a time-structured review of evidence. Journal of Biogeography, 38(5), 817–827. doi: 10.1111/j.1365-2699.2010.02456.x

Pope, L. C., Liggins, L., Keyse, J., Carvalho, S. B., & Riginos, C. (2015). Not the time or the place: The missing spatio-temporal link in publicly available genetic data. Molecular Ecology, 24(15), 3802–3809. doi: 10.1111/mec.13254

Powell, R., Conant, R., & Collins, J. T. (2016). Peterson Field Guide to Reptiles and Amphibians of Eastern and Central North America 4th Edition (4th Editio). Boston, MA: Houghton Mifflin Harcourt.

R Core Team. (2019). R: A language and environment for statistical computing. R Foundation for Statistical Computing, Vienna, Austria. URL https://www.R-project.org/. Vienna, Austria: R Foundation for Statistical Computing.

Radomski, T. P., Hantak, M. M., Brown, A. D., & Kuchta, S. R. (2020). Multilocus phylogeography of eastern red-backed salamanders: cryptic appalachian diversity and postglacial range expansion. Herpetologica, 76(1), 61–73.

Reichman, O. J., Jones, M. B., & Schildhauer, M. P. (2011). Challenges and opportunities of open data in ecology. Science, 331(6018), 703–705. doi: 10.1126/science.1197962

Revell, L. J. (2012). phytools: an R package for phylogenetic comparative biology (and other things). Methods in Ecology and Evolution, 3(2), 217–223. doi: 10.1111/j.2041-210X.2011.00169.x

Rodríguez, A., Börner, M., Pabijan, M., Gehara, M., Haddad, C. F. B., & Vences, M. (2015). Genetic divergence in tropical anurans: deeper phylogeographic structure in forest specialists and in topographically complex regions. Evolutionary Ecology, 29(5), 765–785. doi: 10.1007/s10682-015-9774-7

Romiguier, J., Gayral, P., Ballenghien, M., Bernard, A., Cahais, V., Chenuil, A., … Galtier, N. (2014). Comparative population genomics in animals uncovers the determinants of genetic diversity. Nature, 515(7526), 261–263. doi: 10.1038/nature13685

Semlitsch, R. D. (2008). Differentiating Migration and Dispersal Processes for Pond-Breeding Amphibians. Journal of Wildlife Management, 72(1), 260–267. doi: 10.2193/2007-082

Sexton, J. P., Hangartner, S. B., & Hoffmann, A. A. (2014). Genetic isolation by environment or distance: Which pattern of gene flow is most common? Evolution, 68(1), 1–15. doi: 10.1111/evo.12258

Sidlauskas, B., Ganapathy, G., Hazkani-Covo, E., Jenkins, K. P., Lapp, H., McCall, L. W., … Kidd, D. M. (2010). linking big: The continuing promise of evolutionary synthesis. Evolution, 64(4), 871–880. doi: 10.1111/j.1558-5646.2009.00892.x

Singhal, S., Huang, H., Title, P. O., Donnellan, S. C., Holmes, I., & Rabosky, D. L. (2017). Genetic diversity is largely unpredictable but scales with museum occurrences in a species-rich clade of Australian lizards. Proceedings of the Royal Society B: Biological Sciences, 284(1854). doi: 10.1098/rspb.2016.2588

Smith, B. T., Seeholzer, G. F., Harvey, M. G., Cuervo, A. M., & Brumfield, R. T. (2017). A latitudinal phylogeographic diversity gradient in birds. PLoS Biology, 15(4), 1–25. doi: 10.1371/journal.pbio.2001073

Soltis, D. E., Morris, A. B., McLachlan, J. S., Manos, P. S., & Soltis, P. S. (2006). Comparative phylogeography of unglaciated eastern North America. Molecular Ecology, 15(14), 4261–4293. doi: 10.1111/j.1365-294X.2006.03061.x

Soltis, D. E., & Soltis, P. S. (2016). Mobilizing and integrating big data in studies of spatial and phylogenetic patterns of biodiversity. Plant Diversity, 38(6), 264–270. doi: 10.1016/j.pld.2016.12.001

Stebbins, R. C., & McGinnis, S. M. (2018). Peterson Field Guide to Western Reptiles and Amphibians 4th Edition (4th Editio). Boston, MA: Houghton Mifflin Harcourt.

Stuart, S. N., Chanson, J. S., Cox, N. A., Young, B. E., Rodrigues, A. S. L., Fischman, D. L., & Waller, R. W. (2004). Status and trends of amphibian declines and extinctions worldwide. Science, 306(October), 1783–1786. doi: 10.1126/science.1103538

Thuiller, W., Georges, D., Engler, R., & Breiner, F. (2019). biomod2: Ensemble Platform for Species Distribution Modeling. R package version 3.3-7.1. https://CRAN.R-project.org/package=biomod2.

Vences, M., Thomas, M., Bonett, R. M., & Vieites, D. R. (2005). Deciphering amphibian diversity through DNA barcoding: chances and challenges. Philosophical Transactions of the Royal Society of London. Series B, Biological Sciences, 360(1462), 1859–1868. doi: 10.1098/rstb.2005.1717

Vitt, L. J., Caldwell, J. P., Wilbur, H. M., & Smith, D. C. (1990). Amphibians as harbingers of decay. BioScience, 40(6), 418.

Wang, I. J. (2012). Environmental and topographic variables shape genetic structure and effective population sizes in the endangered Yosemite toad. Diversity and Distributions, 18(10), 1033–1041. doi: 10.1111/j.1472-4642.2012.00897.x

Welsh, H. H., & Ollivier, L. M. (1998). Stream amphibians as indicators of ecosystem stress: A case study from California’s redwoods. Ecological Applications, 8(4), 1118–1132. doi: 10.1890/1051-0761(1998)008[1118:SAAIOE]2.0.CO;2

Whiles, M. R., Lips, K. L., Pringle, C. M., Kilham, S. S., Bixby, R. J., Brenes, R., … Peterson, S. (2006). The effects of amphibian population declines on the structure and function of Neotropical stream ecosystems. Frontiers in Ecology and the Environment, 4(1), 27–34. doi: 10.1017/CBO9781107415324.004

White, E. P., Ernest, S. K. M., Kerkhoff, A. J., & Enquist, B. J. (2007). Relationships between body size and abundance in ecology. Trends in Ecology and Evolution, 22(6), 323–330. doi: 10.1016/j.tree.2007.03.007

Wright, S. (1943). Isolation by Distance. Genetics, 28(2), 114–138. doi: Article

Zipkin, E. F., DiRenzo, G. V., Ray, J. M., Rossman, S., & Lips, K. R. (2020). Tropical snake diversity collapses after widespread amphibian loss. Science, 367(6479), 814–816. doi: 10.1126/science.aay5733

