## Supporting Information for "Predicting amphibian intraspecific diversity with machine learning: Challenges and prospects for integrating traits, geography, and genetic data"

**Methods**

*Random forest classification*

We conducted an example of random forest classification to verify that there was sufficient signal in the dataset using a relatively simple binary problem. We aimed to determine whether conservation status of Nearctic amphibian species can be predicted by range characteristics or species traits and which predictor variables are most important. IUCN Red List categories for each species were obtained from AmphibiaWeb and include Least Concern (LC), Near Threatened (NT), Vulnerable (VU), Endangered (EN), Critically Endangered (CR), Extinct in the Wild (EW), Extinct (EX), or Data Deficient (DD). Given the relatively small number of species in each category, we excluded DD species and combined all categories other than LC into a single category (‘nonLC’). We then applied random forest classification using the R package ‘randomForest’ (Liaw & Wiener, 2002). To determine the importance of each predictor variable, we examined the mean decrease in accuracy (MDA) of prediction when a given predictor is randomly permuted while holding all others constant. Downsampling was used to account for the unevenness of the response variable categories, subsampling the majority class (‘LC’) 100 times without replacement to match the sample size of the minority class (‘nonLC’). We built each random forest with 2000 trees, averaged the MDA for each variable across iterations, and assessed model accuracy using classification error rates. After conducting analyses with all predictor variables of interest, we built classifiers with reduced sets of predictors for simplicity of presentation. We examined whether the most important variables were significantly different for ‘LC’ and ‘nonLC’ species using Student’s t-tests in R.

**Results**

*Predictors of IUCN Conservation Risk*

IUCN conservation status has been assigned for 253 species, with 169 species ranked as “Least Concern” and 84 species in one of the other risk categories. For the initial model of IUCN status with 253 amphibian species, the out-of-bag (OOB) estimate of error rate for the random forest classifier including all variables of interest was 17.8%, with a higher classification error for the ‘nonLC’ class (31.0%) compared to the ‘LC’ class (11.2%). The OOB error rate for a model with a subset of predictors was 15.4% overall, with 30.0% for ‘nonLC’ and 8.3% for ‘LC’. Using downsampling, average error rates were 20.1% (overall) and error rates for the two classes were more similar: 19.8% (nonLC), and 20.5% (LC). For the dataset with a subset of predictors, the average error rates were 19.6% (overall), 17.0% (nonLC), and 22.2% (LC).

The most important predictors of IUCN status were consistently total range size and latitudinal extent (Fig. S5a), regardless of which predictor set was used and whether or not downsampling was employed. Other important predictors were the number of GBIF occurrence records for a species and the standard deviation of bioclimatic variables including temperature seasonality (bio4), precipitation of the driest month (bio14), minimum temperature of the coldest month (bio6), and mean temperature of the driest quarter (bio9). Species traits were not predictive of IUCN status for this dataset. As expected given the criteria used for IUCN risk assessments, which incorporate aspects of population size and geographic range, the species ranked as Least Concern had larger ranges (Fig. S5b; t = 8.14, p = 7.9 x 10^-14^), broader latitudinal extents (Fig. S5c; t = 11.43, p < 2.2 x 10^-16^), more GBIF occurrence records (Fig. S5d; t = 5.80, p = 2.9 x 10^-8^), and more variation in bioclimatic variables (e.g., bio4: Fig. S5e; t = 9.68, p < 2.2 x 10^-16^) than non-LC species.

**Table S1**. Phylogenetic signal values for continuous traits in amphibians. Pagel’s lambda was estimated using the phylosig() function in Phytools (Revell, 2012). Phylogenetic relationships were obtained from VertLife.org (Jetz & Pyron, 2018).

| **Trait** | **Dataset** | **N species** | **Lambda** | **P-value** |
| --- | --- | --- | --- | --- |
| Body size | Amphibians | 285 | 0.999 | 5.62e-73 |
|  | Amphibians with *cytb* | 137 | 0.982 | 1.79e-29 |
|  | Caudata with *cytb* | 98 | 0.991 | 3.14e-21 |
| Larval period min | Amphibians | 285 | 0.940 | 1.15e-47 |
|  | Amphibians with *cytb* | 137 | 0.959 | 3.13e-19 |
|  | Caudata with *cytb* | 98 | 1.000 | 4.53e-17 |
| Larval period max | Amphibians | 285 | 0.976 | 3.10e-37 |
|  | Amphibians with *cytb* | 137 | 0.883 | 8.77e-11 |
|  | Caudata with *cytb* | 98 | 1.000 | 3.66e-13 |
| Clutch size min | Amphibians | 285 | 0.370 | 2.53e-14 |
|  | Amphibians with *cytb* | 137 | 0.608 | 4.02e-06 |
|  | Caudata with *cytb* | 98 | 0.756 | 2.05e-13 |
| Clutch size max | Amphibians | 285 | 0.353 | 7.92e-14 |
|  | Amphibians with *cytb* | 137 | 0.531 | 9.00e-07 |
|  | Caudata with *cytb* | 98 | 0.213 | 0.0144 |

**Figure S1**. List of traits compiled for Nearctic amphibian species and the number of species with available data for each trait. Data were obtained primarily from AmphiBIO (Oliveira, São-Pedro, Santos-Barrera, Penone, & Costa, 2017), species accounts in (AmphibiaWeb, 2018), information from field guides (Powell, Conant, & Collins, 2016; Stebbins & McGinnis, 2018) and primary literature (Bonett, Steffen, Lambert, Wiens, & Chippindale, 2014). Full datasets used for analyses are available from Dryad.

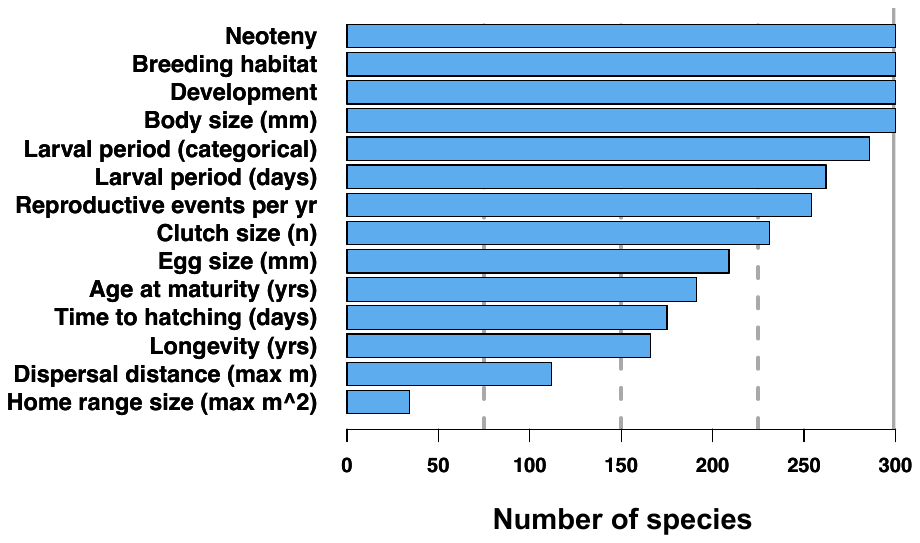

**Figure S2**. Number of amphibian species with sequence alignments for the top 10 genes available on GenBank. Mitochondrial (mito) genes shown in light purple, nuclear genes in dark purple.

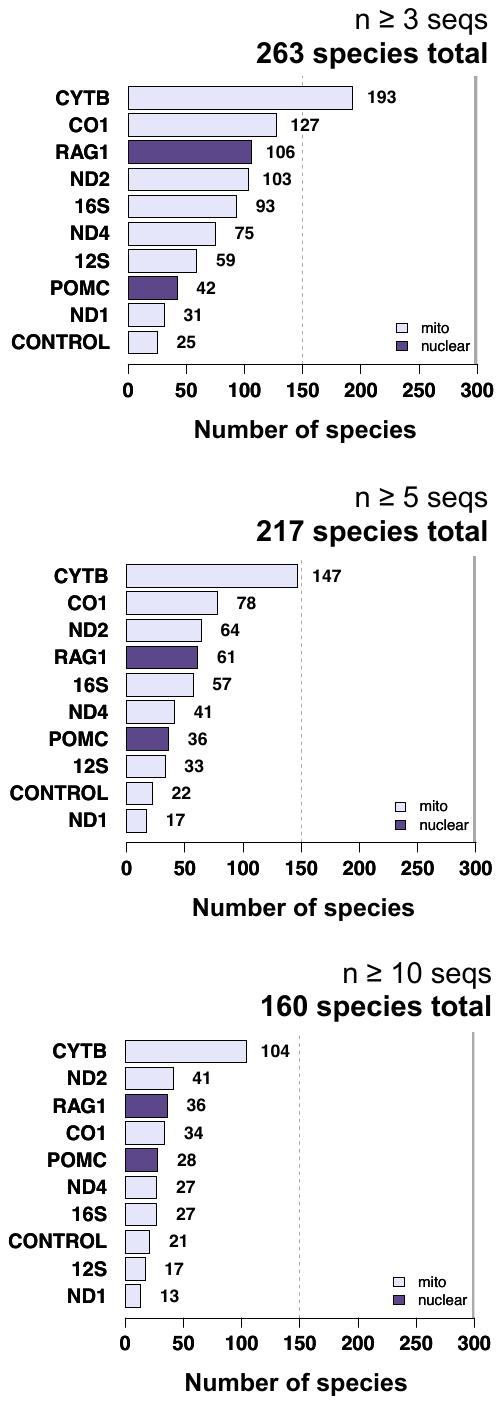

**Figure S3**. Examples demonstrating the variance in nucleotide diversity estimates from 100 datasets of randomly sampled sequences for each sample size from two to 25 sequences. Purple dots represent each replicate dataset, triangles represent median nucleotide diversity from 100 datasets, and the dotted line shows the original nucleotide diversity from the full *cytb* dataset without random subsampling. With >5 sequences included in the alignments, variance in nucleotide diversity estimates decreases rapidly. The same graphs for all species are provided on Dryad. Photos by LNB.

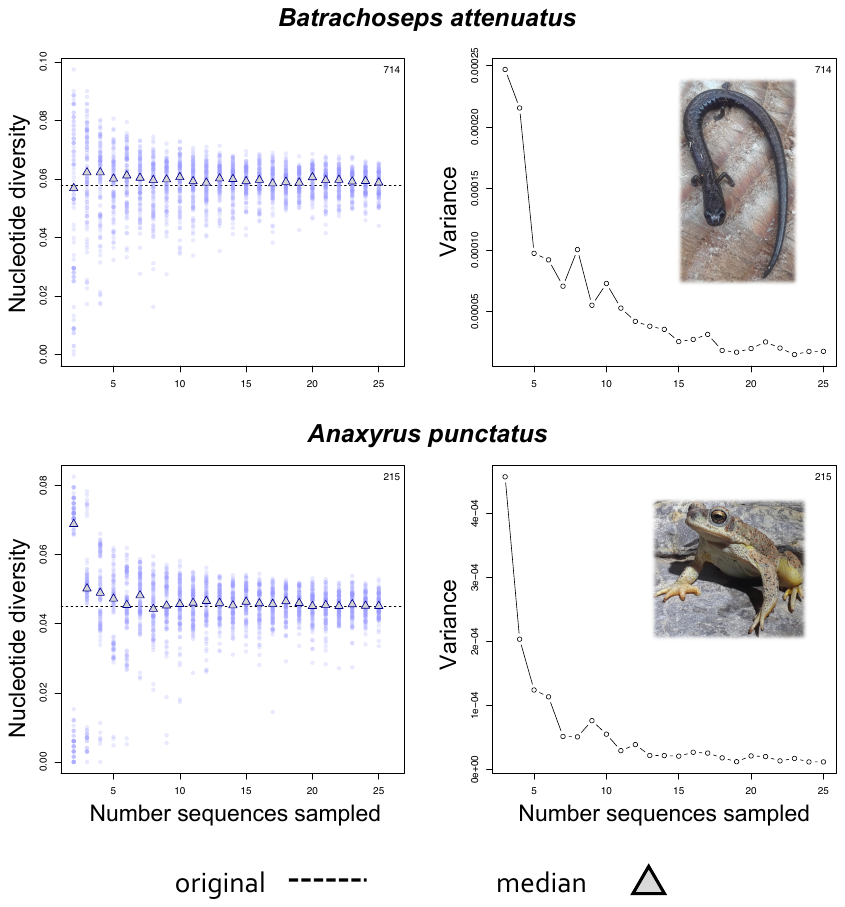

**Figure S4**. Minimum latitude and cytb nucleotide diversity for 98 Caudata species (a,b) and 67 Caudata species (c,d). Nucleotide diversity has a negative relationship with the minimum latitude of a species range. (a,c) The median nucleotide diversity from 100 datasets with five randomly-sampled sequences is shown. (b,d) Phylogenetic independent contrasts for the same metrics using the pruned phylogeny from VertLife.org (Jetz and Pyron 2018).

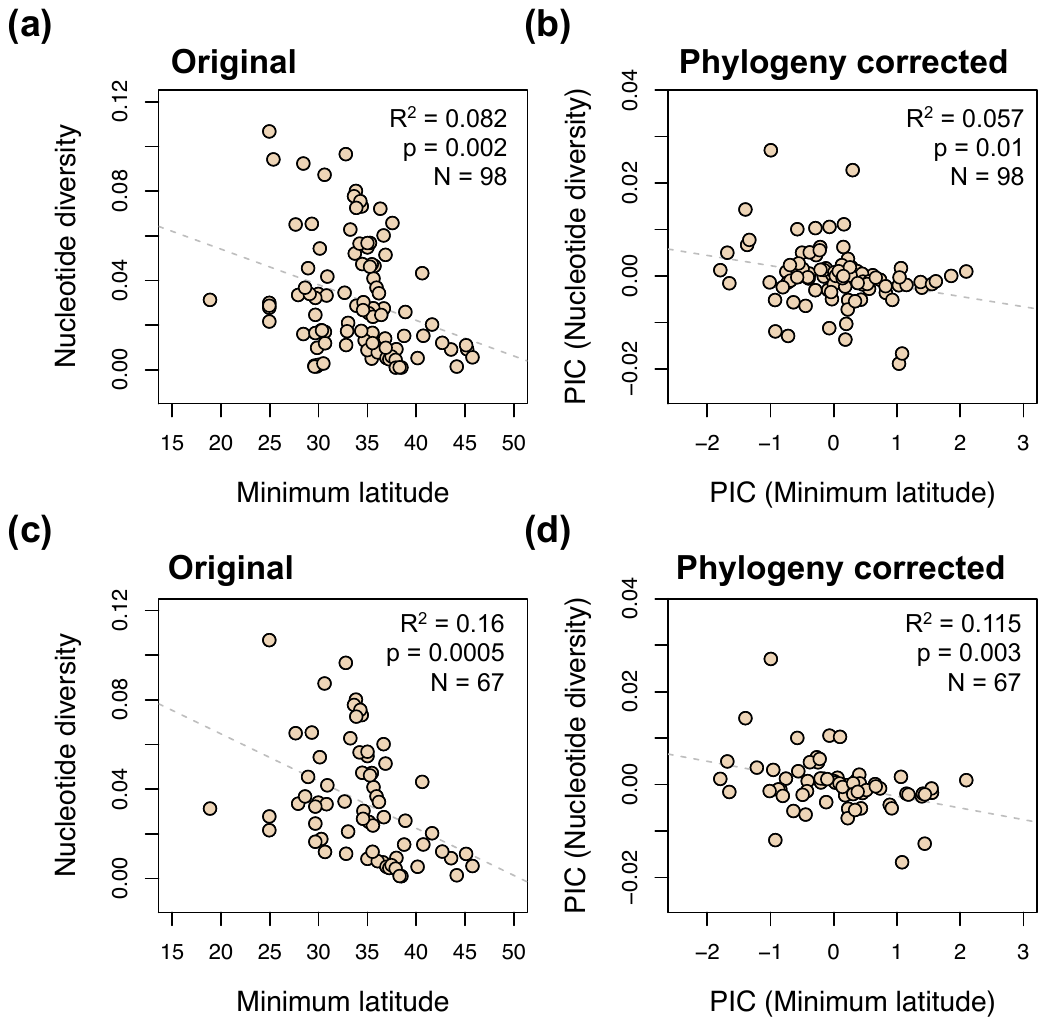

**Figure S5**. Predictors of IUCN status for Nearctic amphibians. a) Average variable importance (mean decrease in accuracy) for each predictor from 100 iterations is shown. The Least Concern (LC) class was downsampled to match the sample size of non-Least Concern (nonLC) species. b-e) Comparison of LC and nonLC species for the top four predictors. As expected based on the criteria used for IUCN status rankings, LC species have larger ranges and associated variables compared to nonLC species.

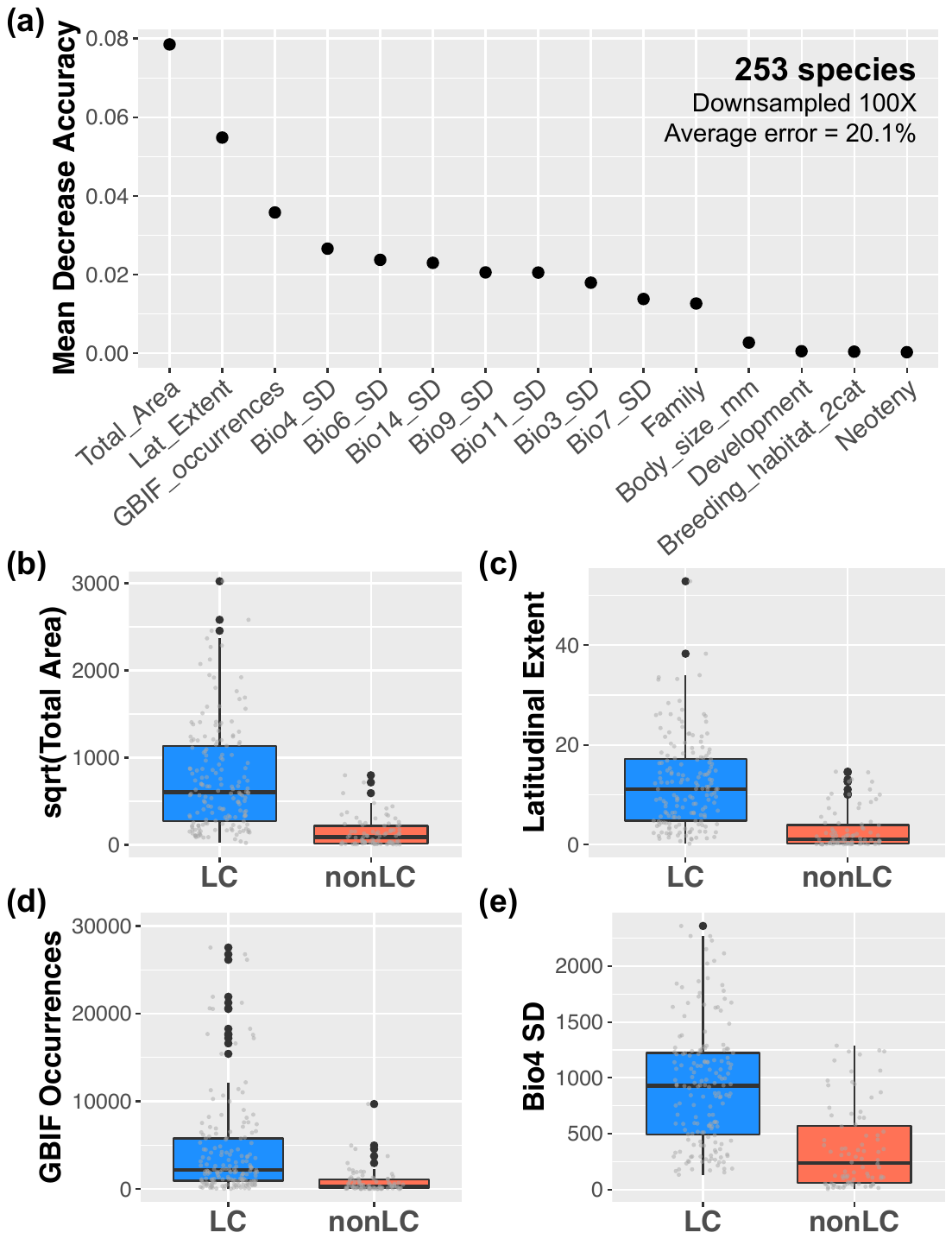

**Literature Cited**

AmphibiaWeb. (2018). <https://amphibiaweb.org>. Retrieved December 18, 2018, from University of California, Berkeley, CA, USA. Accessed 18 Dec 2018. website: https://amphibiaweb.org

Bonett, R. M., Steffen, M. A., Lambert, S. M., Wiens, J. J., & Chippindale, P. T. (2014). Evolution of paedomorphosis in plethodontid salamanders: Ecological correlates and re-evolution of metamorphosis. *Evolution*, *68*(2), 466–482. doi: 10.1111/evo.12274

Jetz, W., & Pyron, R. A. (2018). The interplay of past diversification and evolutionary isolation with present imperilment across the amphibian tree of life. *Nature Ecology and Evolution*, *2*(5), 850–858. doi: 10.1038/s41559-018-0515-5

Liaw, A., & Wiener, M. (2002). Classification and Regression by randomForest. *R News*, *2*(3), 18–22.

Oliveira, B. F., São-Pedro, V. A., Santos-Barrera, G., Penone, C., & Costa, G. C. (2017). AmphiBIO, a global database for amphibian ecological traits. *Scientific Data*, *4*, 1–7. doi: 10.1038/sdata.2017.123

Powell, R., Conant, R., & Collins, J. T. (2016). *Peterson Field Guide to Reptiles and Amphibians of Eastern and Central North America 4th Edition* (4th Editio). Boston, MA: Houghton Mifflin Harcourt.

Revell, L. J. (2012). phytools: an R package for phylogenetic comparative biology (and other things). *Methods in Ecology and Evolution*, *3*(2), 217–223. doi: 10.1111/j.2041-210X.2011.00169.x

Stebbins, R. C., & McGinnis, S. M. (2018). *Peterson Field Guide to Western Reptiles and Amphibians 4th Edition* (4th Editio). Boston, MA: Houghton Mifflin Harcourt.
